# Consistent typing of plasmids with the mge-cluster pipeline

**DOI:** 10.1101/2022.12.16.520696

**Authors:** Sergio Arredondo-Alonso, Rebecca A. Gladstone, Anna K. Pöntinen, João A. Gama, Anita C. Schürch, Val F. Lanza, Pål Jarle Johnsen, Ørjan Samuelsen, Gerry Tonkin-Hill, Jukka Corander

## Abstract

Extrachromosomal elements of bacterial cells such as plasmids are notorious for their importance in evolution and adaptation to changing ecology. However, high-resolution population-wide analysis of plasmids has only become accessible recently with the advent of scalable long-read sequencing technology. Current typing methods for the classification of plasmids remain limited in their scope which motivated us to develop a computationally efficient approach to simultaneously recognize novel types and classify plasmids into previously identified groups. Our method can easily handle thousands of input sequences which are compressed using a unitig representation in a de Bruijn graph. We provide an intuitive visualization, classification and clustering scheme that users can explore interactively. This provides a framework that can be easily distributed and replicated, enabling a consistent labelling of plasmids across past, present, and future sequence collections. We illustrate the attractive features of our approach by the analysis of population-wide plasmid data from the opportunistic pathogen *Escherichia coli* and the distribution of the colistin resistance gene *mcr-1*.*1* in the plasmid population.

## Introduction

Bacteria can exchange genetic material via Horizontal Gene Transfer (HGT) mediated by Mobile Genetic Elements (MGEs) such as temperate phages and plasmids. Plasmids act as key vehicles for the dissemination of important traits such as antimicrobial resistance (AMR) and virulence both within and between species (1, 2). The introduction and broad implementation of long-read sequencing for the assembly of bacterial genomes have led to a dramatic increase in the number of complete plasmid sequences (3).

Clustering and classifying complete plasmid sequences into meaningful groups is a crucial step to understanding the epidemiology of plasmid-encoded genes (4). Without a consistent plasmid typing scheme, it is challenging to examine, for example, whether AMR genes are disseminated by a single or several plasmid types, or if particular plasmid types are overrepresented in successful bacterial clones. Current plasmid typing tools struggle to account for the extreme modularity observed in plasmids, where large genomic blocks can be rapidly gained or lost. Traditionally, plasmids have been classified according to their replicon and associated incompatibility (Inc) groups using tools such as PlasmidFinder (5, 6). However, replicon-based typing suffers from the presence of multiple replicons within the same sequence, offers a limited resolution for epidemiological purposes (4) and is only well-established in particular bacteria phyla (e.g. Proteobacteria). Another strategy consists of typing plasmids based on their relaxase, a protein involved in plasmid mobilisation (7, 8), which is in turn limited to plasmids transmissible by conjugation.

Network analyses based on k-mers or average nucleotide identities (ANI) have been proposed as an alternative classification framework (9, 10). This strategy was implemented in the recent release of COPLA, a novel tool to classify sequences into discrete plasmid taxonomic units (PTUs) based on ANI distances and hierarchical stochastic block modelling (11). MOB-suite is another tool that classifies sequences but relies on k-mers observed in the entire plasmid (12, 13). MOB-suite uses Mash distances coupled with complete linkage clustering to partition plasmid sequences by maximising consistency with replicon and relaxase schemes. The use of COPLA is mainly restricted to typing small sets of sequences due to its computation-intensive algorithm while MOB-suite is more scalable. MOB-suite uses a single Mash threshold to cluster plasmid sequences into discrete groups and can fail to accurately cluster collections of MGEs with different sequence sizes or gene gain/loss rates.

Here, we present *mge-cluster*, a novel approach to consistently type and classify MGE. Mge-cluster provides a classification framework that allows for the typing of thousands of input sequences with a runtime faster than existing algorithms and moderate memory usage. Furthermore, in the light of new MGE data, it offers an option to type these new sequences with an existing mge-cluster model and avoids the need to reanalyse previously typed sequences. Mge-cluster considers the entire sequence content by extracting the unitig sequences which are extended nodes (k-mers) in a compressed de Bruijn graph. The presence/absence of unitigs is embedded into a 2D-representation using openTSNE (14–16), a non-linear dimensionality reduction algorithm that permits the addition of new points to an existing embedding. The non-linear aspect of the tSNE algorithm allows for plasmid clusters to be identified at multiple scales of genetic variation. The HDBSCAN clustering algorithm is then finally used to define plasmid clusters in the resulting 2D embedding (17).

We demonstrate the features of mge-cluster by generating a plasmid classification framework for the opportunistic pathogen *Escherichia coli*, one of the leading causes of bloodstream and urinary tract infections globally with a large number of complete plasmid sequences available. In this organism, virulence factors are usually associated with plasmids, which drive the virulence of enteroinvasive, enteropathogenic, enterohemorrhagic, enteroaggregative, and extraintestinal pathogenic *E. coli* (18, 19). Moreover, plasmids are key hosts for AMR determinants such as extended-spectrum β-lactamases and mobile colistin resistance genes contributing to the emergence of *E. coli* multi-drug resistant infections.

Overall, mge-cluster provides a fast and consistent classification framework for MGEs that can be easily distributed to enhance the analysis and tracking of these elements.

## Results

### Test case: Generating a typing scheme for *Escherichia coli* plasmids

To evaluate the applicability and robustness of mge-cluster, we generated a plasmid typing scheme for *E*.*coli* plasmids. We considered all plasmids from the curated PLSDB plasmid database (20, 21) that includes samples from distinct isolation sources, hosts and countries. This dataset contained highly similar sequences that could lead to an overestimation of the performance of mge-cluster. Thus, redundant sequences were filtered using cd-hit-est (see Methods) to select a single representative plasmid among highly similar sequences (n = 6,185). The discarded plasmid sequences (n = 675) were used as a further test set for benchmarking the runtime and memory required for mge-cluster.

After removing uninformative unitigs (k=31) with low variance (0.01, n=680,491), mge-cluster considered 211,198 unitigs as input to generate the classification framework. The resulting unitigs had an average size of 37.52 bp (median=33.00 bp). This left 189 plasmids (3.1%, 189/6,185) without unitigs and these plasmids were excluded from subsequent clustering analysis resulting in 5,996 remaining plasmids in the analysis. The filtered unitig presence/absence matrix was embedded with openTSNE (perplexity=100) and clusters were called using HDBSCAN (min_cluster=30). In total, we obtained 41 discrete plasmid clusters grouping 4,784 sequences (79.8%, 4,784/5,996) with 1,212 sequences remaining unassigned (20.2%, 1,212/5,996) (Figure 1, Supplementary Table S1).

**Figure 1.**
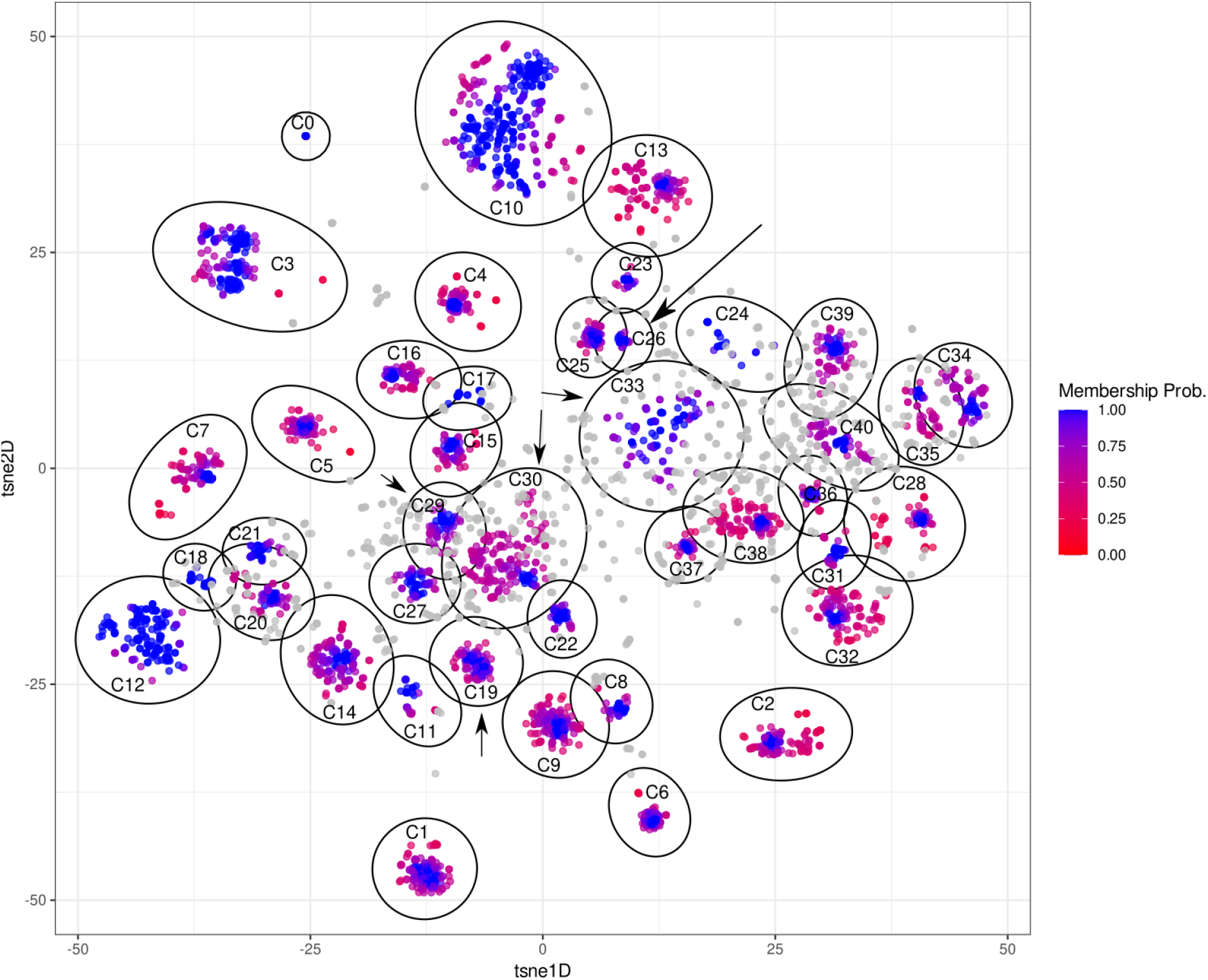
OpenTSNE embedding based on unitigs (k=31) of 5,996 *E. coli* plasmids. Each point corresponds to a plasmid sequence and their assigned cluster (C) is labelled based on the cluster ID (n=41) defined by HDBSCAN. Sequences belonging to an HDBSCAN cluster are coloured (from red to blue) based on their membership probability. Unassigned sequences correspond to plasmids with a membership probability of 0 of belonging to any defined cluster and are coloured in grey. The ellipses (in black) delimit the cluster coordinates and were estimated using the Khachiyan algorithm implemented in the ggforce R package. To facilitate finding clusters 19, 26, 29, 30 and 33, which are highlighted as examples in the text, we indicated their positions with an arrow in the plot.

The chosen perplexity value can impact the non-linear resultant embedding such that low perplexity values tend to preserve the local structure better, while sometimes artificially introducing some structure when none exists. Conversely, high perplexity values tend to preserve more of the global structure at the cost of merging small clusters together. We evaluated the impact of varying this mge-cluster parameter (perplexity=10, 30, 50, 200) by comparing their resulting clustering assignments using the adjusted Rand index. This index can vary from 0 (completely distinct typing models) to 1 (identical typing models) while adjusting for randomly assigning two sequences belonging to the same cluster. We observed that mge-cluster produced assignments robustly (average Rand index=0.95) to the chosen perplexity values when considering sequences assigned by two resulting models (Table 1).

**Table 1.**
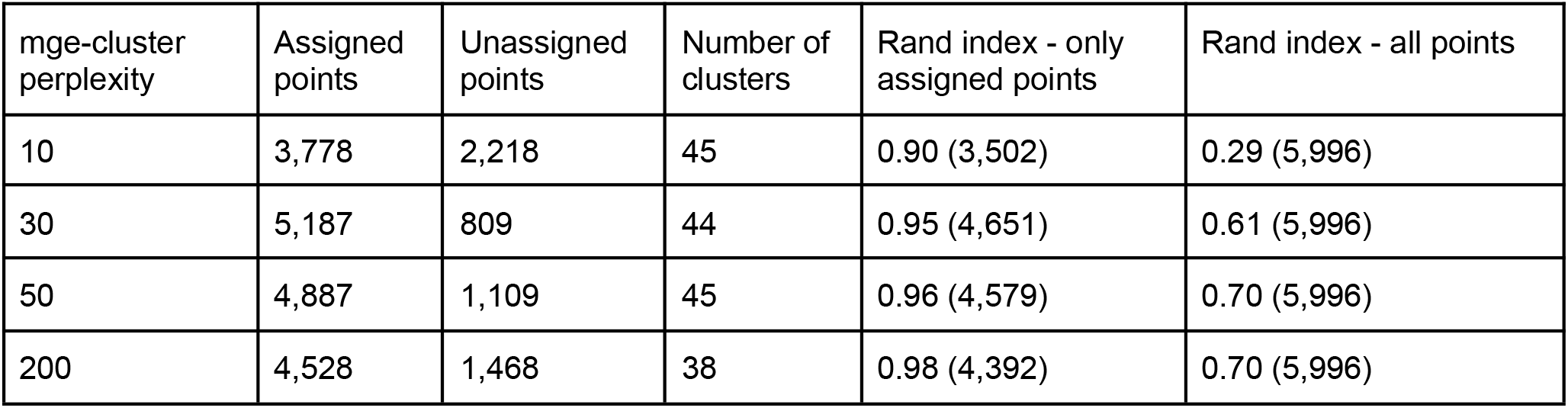
Comparison of the mge-cluster models over a range of perplexity values (10, 30, 50, 200). The models were compared against the mge-cluster solution, corresponding to a perplexity value of 100. The Rand index was first computed considering only points assigned to a cluster by the two clustering solutions and thus ignoring points which were either unassigned by one of the two models. Secondly, the Rand index was computed with all points (assigned and unassigned) to highlight the discrepancy between the models is mainly caused by sequences clustered by one of the two models but unassigned by the other.

| mge-cluster perplexity | Assigned points | Unassigned points | Number of clusters | Rand index - only assigned points | Rand index - all points |
| --- | --- | --- | --- | --- | --- |
| 10 | 3,778 | 2,218 | 45 | 0.90 (3,502) | 0.29 (5,996) |
| 30 | 5,187 | 809 | 44 | 0.95 (4,651) | 0.61 (5,996) |
| 50 | 4,887 | 1,109 | 45 | 0.96 (4,579) | 0.70 (5,996) |
| 200 | 4,528 | 1,468 | 38 | 0.98 (4,392) | 0.70 (5,996) |

In addition, we show that the mge-cluster discrepancy between models can be explained by the sequences which are unassigned by one of the models but clustered in the other (Table 1). Consequently, we encourage users to run mge-cluster by setting distinct perplexity values to evaluate cluster stability.

Plasmids can rapidly incorporate or lose genomic modules or even co-integrate with other sequences present in the same cell, which drastically affects their size. For each cluster (n=41) (perplexity=100, min_cluster=30), the interquartile range (IQR) of the sequence length was on average 18.66 kbp but with pronounced differences depending on the cluster (Supplementary Table S2). As an example, cluster 26 (Figure 1) with a mean length of 94.6 kbp showed an IQR of 0.26 kbp indicating an almost intact plasmid backbone, while, cluster 19 (Figure 1) with a mean length of 159.4 kbp had an IQR of 51.2 kbp indicating the presence of distinct gained/lost genomic modules shared by only a fraction of the plasmids assigned to this cluster.

To quantify the percentage of shared sequences among plasmids from the same cluster, we used *pyani* to retrieve average nucleotide identity (ANI) and coverage values (22). On average, plasmids shared 62.3% of their sequence (pyani coverage) with other members from the same cluster with an associated ANI of 95.7%. We observed that the average coverage shared between plasmids varied substantially among clusters indicating distinct degrees of plasmid modularity as previously exemplified with the IQR of the sequence length (Supplementary Table S2). Clusters 29, 30 and 33, formed by large plasmids, displayed a low pyani coverage indicating that plasmids from those clusters shared only a minor fraction of their sequence. To further understand the content of each mge-cluster, we visualized the diversity of replicons (Supplementary Figure S1) predicted by the MOB-typer module of MOB-suite (12) based largely on PlasmidFinder (6).

### Comparison of mge-cluster against other plasmid typing tools

To assess the level of concordance with current typing schemes, we compared the mge-cluster results against the gold standard methods for plasmid typing. MOB-suite provides a five-character fixed-length code (2 letters and 3 digits) to identify sequences belonging to the same group (termed ‘primary_cluster_id’) (Supplementary Table 3), while COPLA provides a PTU designation (Supplementary Table 4). However, the CPU time (167 minutes, 22 min wall-clock time) and memory (319.5 Mb) required for COPLA to predict the plasmid type of a single sequence (NZ_CP024805.1) hampered us from predicting the entire *E. coli* dataset of 6,185 plasmids for a full comparison with mge-cluster. However, 695 sequences (11.2%, 695/6,185) from our dataset were typed in the original publication describing COPLA [10] and were further considered in this comparison.

To compare the overall clustering concordance, we considered the adjusted Rand index which fluctuates from 0 (different clusters) to 1 (same clusters). We observed a moderate agreement between mge-cluster and MOB-suite with an index of 0.61, while for COPLA the adjusted Rand index was 0.53 (Figure 2, no threshold). Notably, we observed that by increasing the membership probability threshold of mge-cluster to assign plasmids to particular clusters, we observed a higher level of overlap between the tools reaching a maximum adjusted Rand index value of 0.77 (Figure S2, threshold=0.9).

**Figure 2.**
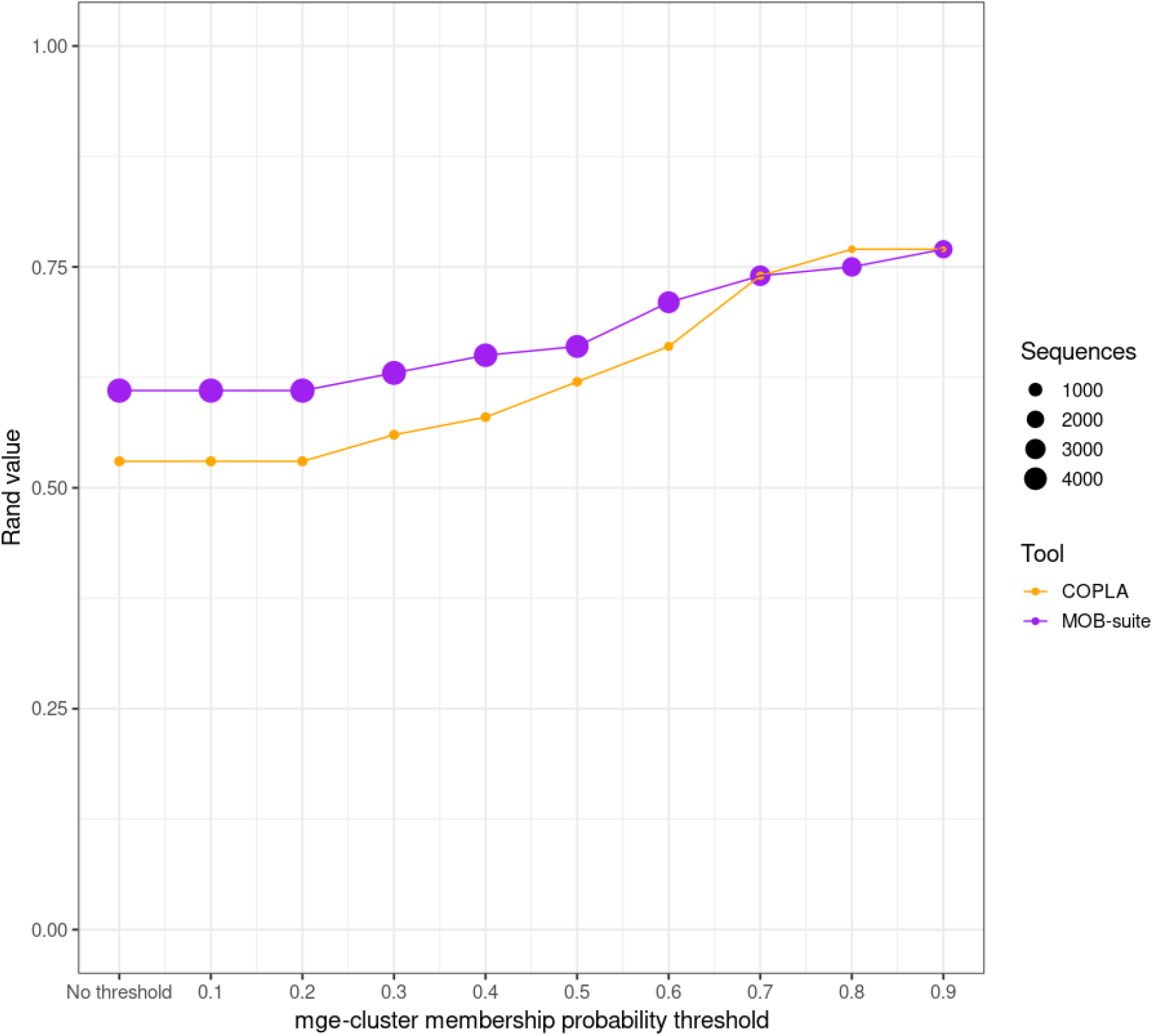
Concordance of mge-cluster results compared to MOB-suite (in purple) and COPLA (in orange) based on adjusted Rand index values. For each sequence assigned to a cluster by mge-cluster, the tool returns a membership probability. This probability was used to set several thresholds (ranging from no threshold to 0.9) to assign the plasmid sequences and assess their concordance against MOB-suite and COPLA. Each point in the comparison is sized according to the number of sequences used to compute the adjusted Rand index value between the tools.

To define which mge-clusters had a higher level of overlap with MOB-suite and COPLA types, we calculated the Simpson diversity of each mge-cluster. For instance, if all plasmids from a particular mge-cluster were designated as a single type by MOB-suite and COPLA, this Simpson diversity value would be 0. In contrast, the presence of multiple types defined by MOB-suite and COPLA would result in diversity values close to 1. The diversity of MOB-suite and COPLA types was represented in Supplementary Figures S2 and S3, respectively.

The overall Simpson diversity per cluster was 0.46 and 0.21 for MOB-suite and COPLA, respectively. We observed that by increasing the membership probability threshold, the average diversity of MOB-suite types was substantially reduced up to 0.23 (threshold=0.9) with no changes in the case of COPLA (0.21, threshold=0.9) (Figure 3). COPLA produced the same PTU designation (PTU-FE) for 10 distinct mge-clusters which resulted in a lower Simpson diversity than for MOB-suite at the cost of merging together plasmids with a distinct core gene content (Supplementary Figure S3). This PTU-FE type was reported in COPLA’s publication as problematic because several plasmid configurations were present resulting in a low intra-cluster density (10).

**Figure 3.**
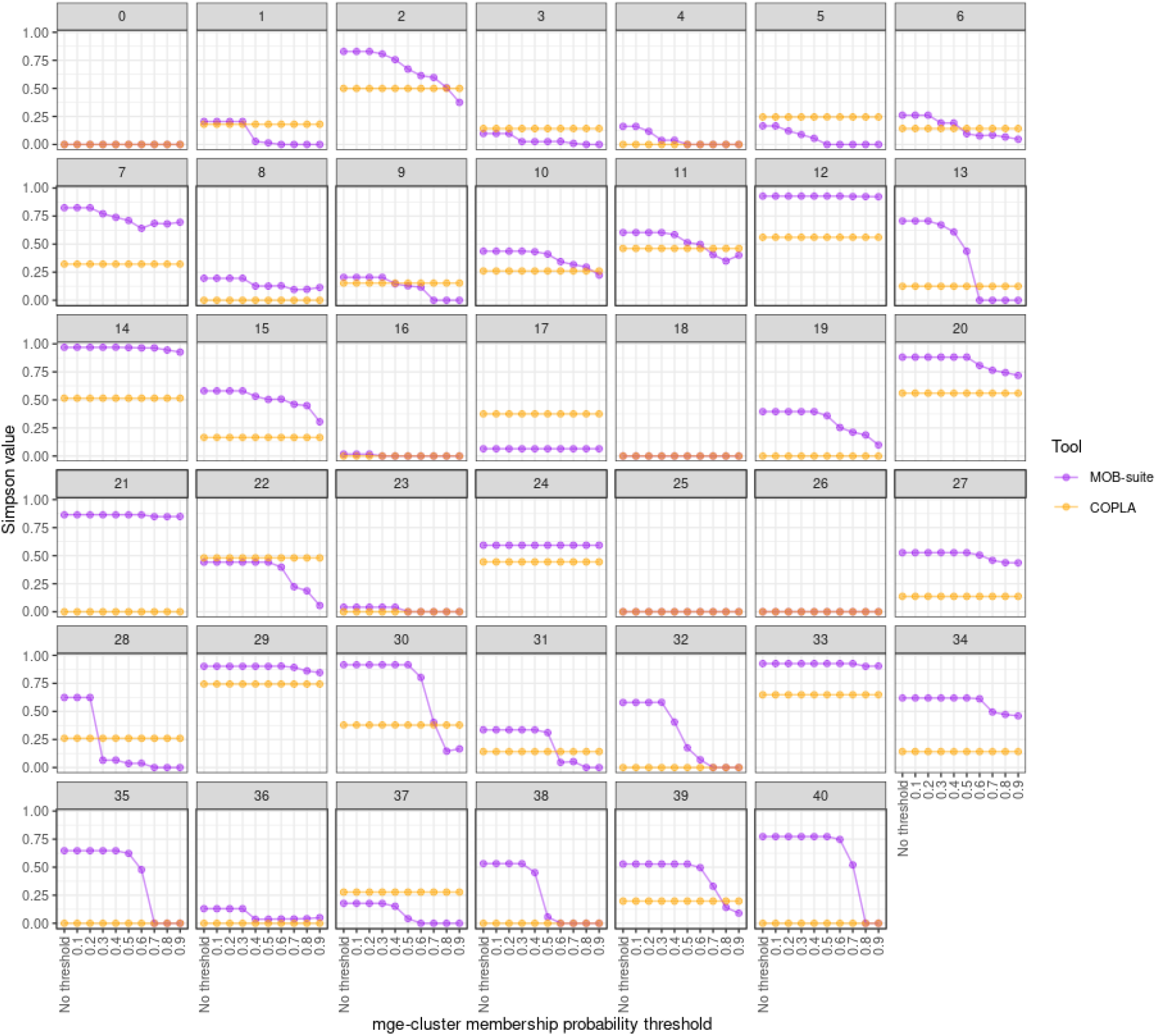
Diversity of MOB-suite (in purple) and COPLA (in orange) types for each mge-cluster (n=41). The diversity (Simpson value) could range from 0 (agreement with mge-cluster) to 1 (no agreement). The probability assigned to each plasmid by mge-cluster was considered to set several thresholds (ranging from no threshold to 0.9) to assign the plasmid sequences and assess their concordance with MOB-suite and COPLA.

MOB-suite showed a consistent agreement (Simpson value < 0.2) in 13 mge-clusters (Figure 3, Supplementary Table S2). The disagreement between both tools occurred in the mge-clusters with an average small plasmid length (< 10 kbp) (clusters: 2, 7, 12, 14, 20, 21). These clusters consisted of sequences with a predominant replicon type (Supplementary Figure S1), however, MOB-suite predicted those sequences in distinct clusters (Supplementary Figure S2). MOB-suite confirmed with a high Simpson diversity value, that clusters 29, 30 and 33 were formed by large plasmids from distinct types (Supplementary Table S2).

For the rest of the clusters, we observed that mob-cluster only tended to group sequences that were highly similar in their gene content (high identity and coverage). To illustrate this, we considered a random sequence from mge-cluster 31 predicted with a different type by MOB-suite (NZ_LT985213.1 for AA735, NZ_CP010138.1 for AA334) and performed a gene synteny analysis (Supplementary Figure S4). We could observe that these two sequences, despite being classified by MOB-suite as distinct types (AA735 and AA334), had a blastn coverage and identity of 73.1% and 99.6%, respectively. The synteny analysis revealed both sequences had an IncFII replicon with a well-conserved synteny (Supplementary Figure S4).

However, NZ_LT985213.1 had incorporated an extra module corresponding to the co-integration of an IncFIA replicon. MOB-suite uses a stringent Mash threshold (0.06) to group plasmid sequences. Therefore, sequences that share a highly similar plasmid backbone but have gained/lost genomic modules or even co-integrated other plasmids tend to be grouped by MOB-suite into distinct types. In the case of mge-cluster, plasmids acquiring an extra genomic module have a lower membership probability of belonging to the cluster since their unitig content differs but are still part of the same cluster. This behaviour explains why the increase in the membership threshold of mge-cluster results in a higher agreement with MOB-suite (Figure 2).

### Predicting novel sequences with an existing mge-cluster model

Mge-cluster was built to generate a classification network that can also assign the same cluster names without the requirement to re-analyze any previous dataset and to keep consistent cluster names (*--existing* mode). We considered the sequences discarded by cd-hit-est (n=675) to benchmark the runtime and memory required by mge-cluster to assign these sequences to the previous clusters. In addition, these sequences should be embedded and assigned to the same cluster as the representative sequence from the cd-hit-est step.

Mge-cluster predicted these 675 samples using less CPU and wall-clock time (23.3 minutes, ∼4 min wall-clock time) than for MOB-suite (CPU time 32.2 minutes, ∼ 26 minutes). However, the peak memory usage of mge-cluster (15.9 Gb) was substantially higher than for MOB-suite (4.5 Gb). From these 675 samples, 15 sequences corresponded to cd-hit clusters for which its representative sequence was discarded in the mge-cluster model because of the absence of unitigs and were not evaluated further. Mge-cluster correctly assigned 99.2% (655/660) of the plasmids to the same cluster as their corresponding reference sequence (Suppl. Figure S5). In five cases (0.8%, 5/660), mge-cluster predicted another cluster, including four cases where the model returned an unassignment (−1) category.

Next, we evaluated the performance of mge-cluster predicting plasmids not present in *E. coli* and thus unseen by the pipeline to build the mge-cluster model. For this, we considered all *Staphylococcus aureus* plasmids (n=1,021) from the PLSDB database because of the absence of plasmid transmission events between these two species (9, 10). Mge-cluster did not detect any *E*.*coli*-specific unitigs (0/211,198) for 972 *S. aureus* plasmids (95%) and thus those sequences were not assigned to any of the mge-clusters from the *E. coli* model (Supplementary Table S5). This is due to the high specificity of the unitigs used in the mge-cluster model which had a minimum size of 31 bp and an average size of 37.52 bp. From the remaining 49 plasmids (5%), 28 plasmids were not assigned to any cluster, 12 plasmids were assigned to the mge-cluster 29 and 8 plasmids to the mge-cluster 30. We confirmed that the plasmids assigned to mge-clusters 29 and 30, had a low number of unitigs present and thus corresponded to samples with a vector of nearly all zeros. In those cases, mge-cluster embedded those sequences into clusters 29 and 30 which we previously highlighted as random noise clusters.

Lastly, we assessed the performance of mge-cluster predicting plasmids likely shared in other bacterial species from the same family (*Enterobacterales*) as *E. coli*. For this, we selected plasmids from the incompatibility group N (IncN) since they have a conserved core genome, which was used to develop a specific plasmid multilocus sequence typing (pMLST) scheme (6) and have been reported across several bacterial species belonging to *Enterobacterales* (23). We considered all IncN non-*E*.*coli* plasmids from the PLSDB database containing uniquely a single replication gene (n=206) and predicted their clustering assignment with the *E. coli* mge-cluster model (Supplementary Table S6). We observed that most IncN plasmids (80.6%, 166/206) were predicted as part of the mge-cluster 27 which contains a majority of *E. coli* plasmids belonging to this incompatibility group (Supplementary Figure 1a) and thus confirming that this plasmid type is shared and has a conserved genomic backbone among *Enterobacterales*. In total, 34 plasmids (16.5%) could not be assigned to any mge-cluster and were labelled as (−1) showing that some of these IncN plasmids might have acquired or recombined with other genomic modules and thus have a clearly distinct unitig content. The remaining plasmids (2.9%, 6/206) were scattered among mge-clusters 29 (n=4), 14 (n=1) and 30 (n=1).

### Cluster distribution and visualization of a gene of interest in the embedding space

The typing scheme offered by mge-cluster is optimal for visualizing the genomes carrying any particular gene of special interest and tracking its distribution in future sequencing studies. To illustrate this, we considered the AMR gene *mcr-1*.*1* which confers resistance to colistin, a last-resort antibiotic for treating infections caused by multi-drug resistant *E. coli*. This AMR gene was first reported in 2016 on a plasmid with an IncI2-type backbone (24) that can be mobilised among distinct MGEs by the presence of an *ISApl1* transposon element situated upstream of the gene (25).

We observed that 327 plasmids contained the *mcr-1*.*1* gene, the vast majority of these present in only three mge-clusters: 3 (n=168), 1 (n=71) and 16 (n=53) (Figure 4). This was in agreement with previous reports (26, 27) showing this AMR gene to be mainly spread by the plasmid backbones IncI2 (mge-cluster 3), IncHI2 (mge-cluster 1) and IncX4 (mge-cluster 16) (Supplementary Figure S1). However, we also observed that the AMR gene was present in nine additional mge-clusters (30/327, 9.2%) (Figure 4) and 5 sequences (1.5%) could not be assigned to any mge-cluster. This illustrates how a consistent typing provided by mge-cluster can be used to explore whether these nine clusters represent spillover events of the gene to other plasmid backbones for which the gene might be further disseminated using new plasmid types.

**Figure 4.**
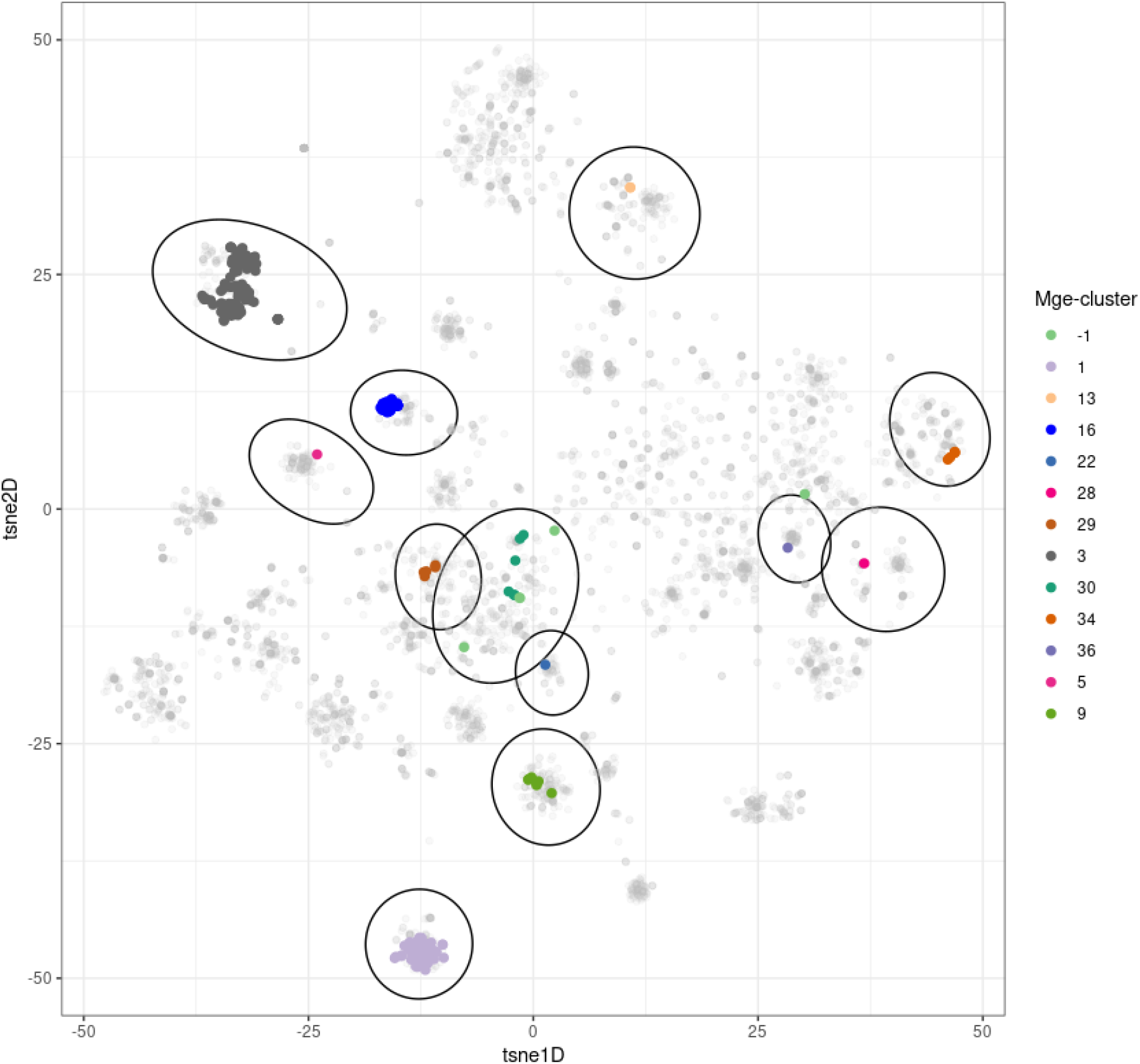
Distribution of the *mcr-1*.*1* gene on the embedding space created by mge-cluster. The plasmids (n=327) containing the gene are coloured in the plot according to their cluster labels. The clusters (n=12) containing at least a single sequence having the *mcr-1*.*1* gene are indicated with an ellipse using the Khachiyan algorithm implemented in the ggforce R package. The sequences which were not assigned to any mge-cluster were labelled as ‘-1’ (light green) in the legend.

## Discussion

The number of MGEs available in public databases has exploded since the introduction of long-read sequencing technologies. However, the contextualization and comparison of MGEs are hampered by their high rates of recombination which result in the absence of a conserved marker that can be broadly used by standard phylogenetic methods. Mge-cluster responds to this need by generating discrete clusters from sequences generally evolving through a fast and dynamic turnover of gene gain/loss events.

We demonstrated the potential of mge-cluster by developing an *E. coli* model to classify plasmid sequences. We observed that the clusters generated by mge-cluster typically consisted of sequences with a shared plasmid backbone (coverage ∼62%) but distinct accessory content. Mge-cluster and MOB-suite showed a moderate level of agreement between clustering solutions. Some of the disagreement between the tools is explained by mge-cluster grouping together plasmid sequences that have acquired an extra replicon sequence due to the cointegration of another plasmid. This characteristic of mge-cluster is particularly beneficial for tracking a plasmid in the context of longitudinal studies for which the same plasmid can rapidly gain/lose genomic modules. The current version of COPLA makes the typing of large collections unfeasible due to the CPU time required to run a single sample. Moreover, in the particular case of *E. coli*, COPLA erroneously merges clusters from distinct plasmids under the PTU-FE group. In contrast to MOB-suite and COPLA, mge-cluster does not require a predefined distance threshold to generate the typing model which facilitates broad applicability across distinct species and datasets.

For epidemiological purposes, clusters obtained with mge-cluster should be interpreted in a similar manner as MLST (28) or BAPS groups (29) that cluster strains based on chromosomal housekeeping gene alleles and genome alignments respectively. Even if two plasmids from different samples belong to the same cluster, we cannot directly assume a plasmid transmission scenario. For this, mge-cluster can be considered as a starting point to perform secondary analyses such as SNP phylogenies based on the resulting cluster core genome. These secondary analyses can confirm or refute transmission links, as recently illustrated in two studies presented by Ludden *et al*. and Hawkey *et al*. (30, 31). These types of sequencing studies may benefit from the usage of mge-cluster to define plasmid discrete groups which opens the possibility of sharing their models with other groups for further tracking the distribution of a plasmid or gene-of-interest.

While we demonstrated the use of mge-cluster using a single species, the pipeline can also be run on more diverse datasets such as the combination of plasmid sequences from the *Enterobacterales* family. To showcase mge-cluster we considered complete plasmid sequences, but the pipeline could also be utilised to type the bins resulting from tools predicting and extracting plasmids from short-read sequencing data (12, 32, 33). We anticipate that mge-cluster can in addition be used for generating discrete clusters from other types of MGEs with sufficient gene content diversity including phages, integrative and conjugative elements (ICEs) or flanking sequences surrounding a gene-of-interest (e.g AMR gene).

The ability of mge-cluster to rapidly assign new plasmids with a consistent type facilitates the comparison of plasmids derived from distinct collections and boosts our capacity to conduct MGE surveillance in general.

## Materials and Methods

### Mge-cluster workflow

Mge-cluster is a Python package installable through bioconda https://anaconda.org/bioconda/mge-cluster, freely available under the open-source MIT license https://gitlab.com/sirarredondo/mge_cluster. Figure 5 illustrates the two different operational modes of mge-cluster: *--create* and *--existing*. In both cases, mge-cluster takes as input a file that indicates the absolute or relative paths to the nucleotide sequence files (.fasta format). The *--create* mode will generate a new classification scheme for the sequences provided as input by the user while the *--existing* mode will return embedding coordinates and cluster assignments considering a previous, existing mge-cluster model. Both modes can be run with the multithreading option (*--threads*) to reduce mge-cluster runtime.

**Figure 5.**
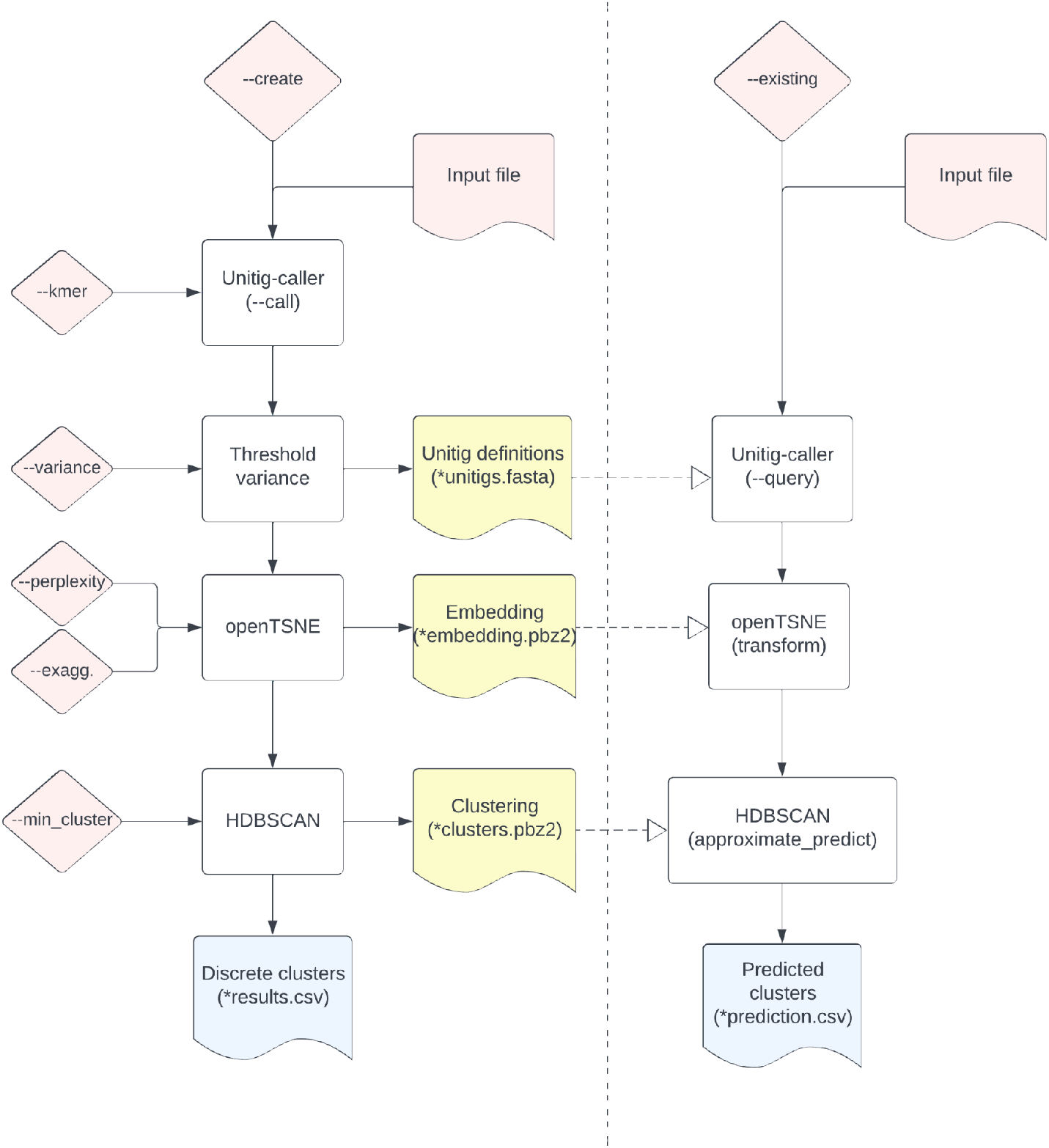
Summary of the mge-cluster workflow. The tool is composed of two distinct operational modes: *--create* (left) and *--existing* (right). Both modes require as input a file listing the absolute or relative paths of the nucleotide sequences. The *--create* mode of mge-cluster requires the following arguments (*--kmer, --variance, --perplexity, --exaggeration* and *--min_cluster*) to generate discrete clusters from the sequences provided in the input. The *--existing* mode of mge-cluster requires the files (in yellow) generated by the *--create* mode to predict the clusters of a new batch of nucleotide sequences.

### Operational mode *--create*: Unitigs as classification features

Unitigs defined as extended nodes in a compressed de Bruijn graph were selected as features for building the classification framework. Unitig-caller (*--call* mode, version 1.2.1) https://github.com/bacpop/unitig-caller which implements Bifrost Build (34) is used with a k-mer size specified by the mge-cluster (argument *--kmer*) to generate a presence/absence matrix of the unitigs present in the input file.

Bifrost initially considers a *de Bruijn* graph structure defined as a direct graph:

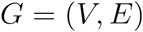

*V* corresponds to the number of vertices (k-mers) present and *E* to the edges connecting the distinct vertices. Thus, the vertices *V* present in graph *G* can be defined by:

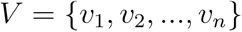

The edges *E* can be defined as direct connections between two vertices of *V*:

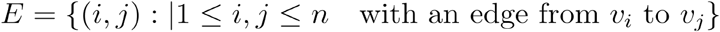

For each *v* ∈ *V*, we define the in-degree *d*^*i*^(*v*) and out-degree *d*^*0*^(*v*) as the number of edges in *E* towards and from *v* respectively.

Paths in the graph can be defined as finite sequences of distinct vertices connected by edges *p* = (*v*_0,_ *e* (*v*_*0*_,v_1_), *v*_1,_ *e*(*v*_1,_ *v*_2_),…,*v*_*k*_,*e*(*v*_*k*−1_*v*_*k*_)). Bifrost then considers all non-branching paths, defined as paths *p* in which all vertices have an *d*^*i*^(*v*) = 1 and *d*^*0*^(*v*) = 1 excluding the first and last vertices in *p*.

Each non-branching path is merged into a single vertex, termed unitig in Bifrost. Those unitigs represent extensions of the initial k-mers (vertices) that are longer in length than the original k-mer size. Unitig-caller then creates a presence/absence matrix of those unitigs. We can define *M*, as a binary matrix with *s* * *u* dimensions, in which *s* is the total number of sequences present in the input file and *u* corresponds to the total number of unitigs extracted by Bifrost.

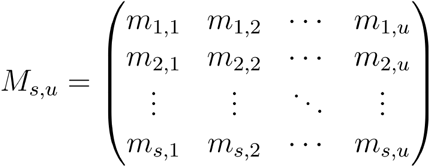

Unitigs were chosen over other features (e.g. gene presence/absence) because identical unitig definitions could be computed between distinct datasets, an essential characteristic for the *--existing* prediction mode of mge-cluster. To reduce the memory use required to build the typing scheme, we remove unitigs with a variance less than 0.01 (default) using the function VarianceThreshold of the python package sklearn (version 1.0.2) (35). In this manner, we remove unitigs (features in the model) that have the same value for all samples and thus do not provide any relevant information for the embedding process. This variance threshold can be modified by the user in mge-cluster (argument *--variance*).

### Operational mode *--create*: Embedding the presence/absence of unitigs into a lower number of dimensions

We considered the implementation of the tSNE algorithm available in the python package openTSNE (version 0.6.1) (15) to generate a 2D embedding based on *M*, the unitigs presence/absence matrix. This new implementation improved the global positioning of the points and introduced the possibility of mapping new points into an existing, reference embedding. The multidimensional presence/absence matrix of unitig-caller can be represented as *M* = {*m*_1,_ *m*_2_,…,*m*_*s*_} ∈ *R*^*u*^ for which *m*_*s*_ corresponds to a datapoint (sequence) with *u* defined as the number of dimensions (number of unitigs passing the variance threshold). In our case, openTSNE is run to find a 2D dimensional embedding *Y* = {*y*_1,_ *y*_2_,…,*y*_*s*_} ∈ *R*^*2*^ in which the original distance between *m*_1_ and *m*_2_ is preserved in *y*_1_ and *y*_2_. The similarity between two data points in the original space is measured with Jaccard distances (flag --metric). The perplexity value is one of the main parameters of openTSNE that affects how the similarity between two data points in the original space is preserved in the resulting embedding space. Large perplexity values tend to preserve the global structure of the data better while obscuring some of the local structure potentially resulting in small clusters being merged together. Small perplexity values generate tight dense clusters preserving the local structure better but ignore the overall global structure for which the distance and position of the clusters in the resulting embedding can no longer be considered.

The TSNE function can be run with different perplexity values specified by the user with the mge-cluster arguments *--perplexity*, using ‘exact’ as the method for finding the nearest neighbor (flag *--neighbors*). For reproducibility purposes, we fixed the seed of the random number generator with the flag *--random_state*.

### *Operational* mode *--create*: Calling plasmid clusters in the embedding space

To define which clusters were present in the embedding space *Y* = {*y*_1,_ *y*_2_,…,*y*_*s*_} ∈ *R*^*2*^ created by openTSNE, we required a clustering algorithm that (i) did not force us to provide the number of clusters present in the data, (ii) tolerated noisy data since plasmid modularity can result in sequences that are hybrids between two neighbouring clusters, (iii) tolerated clusters with different density and sizes (iv) allowed the assignment of new data points to an existing clustering solution. Based on these four premises, we selected the HDBSCAN algorithm (17), an improved version of dbscan that finds highly stable clusters over a range of epsilon values (the main parameter of dbscan).

HDBSCAN defines the mutual reachability distance (extracted from HDBSCAN documentation) as *d*_*mreach−k*_(*y*_1_,*y*_*s*_) = *max* (*core*_*k*_(*y*_1_), *core*_*k*_(*y*_s_), *d*(*y*_1,_*y*_*s*_)), where *d*(*y*_1,_*y*_*s*_) is the original metric distance (Euclidean) and *core*_*k*_ the distance to its *k*th neighbour. This *mreach* distance is used to transform the embedding space into a new space where points with low core distances remain together while pushing away sparser points. This distance is considered to create a graph structure *HG* = (*P,D*) in which nodes *P* correspond to the original data points *y*_*s*_ while *D* are all edges with weight equal to *d*_*mreach−k*_(*y*_1_,*y*_*s*_). HDBSCAN then transforms *HG* into a minimum spanning tree to look into the hierarchy of connected components. Lastly, HDBSCAN uses the parameter --min_cluster_size to define the minimum number of points that are required to define a cluster. This parameter is then used to generate a condensed tree to select clusters with high persistence. Lastly, HDBSCAN outputs for each point their assigned cluster and membership probability.

The python package hdbscan (version 0.8.28) with the primary function HDBSCAN is run to specify a default minimum cluster size (flag *--min_cluster_size*) defined by the user in mge-cluster (argument *--min_cluster*).

The main output of this operational mode consists of a comma-separated file (csv) with the embedding coordinates given by openTSNE (columns ‘tsne1D’, ‘tsne2D’), the cluster assigned and membership probability returned by HDBSCAN (column ‘Standard_Cluster’ and ‘Membership_Probability’ and the last column (‘Sample_Name’) indicating the header extracted from the given nucleotide sequences.

### Operational mode *--create*: Storing and distributing an mge-cluster model

Mge-cluster was specifically designed to generate a classification scheme that can easily be distributed and reused by other users. The following files constitute a mge-cluster model: i) *\**.*unitigs*.*fasta*, the fasta file containing the unitigs with a variance higher than specified in the argument *--variance*, ii) *\**.*embedding*.*pbz2*, embedding model created by openTSNE to transform the unitig presence/absence matrix into 2D and iii) *\*clusters*.*pbz2*, clustering model created by HDBSCAN to call clusters in the resulting embedding from openTSNE.

### Operational mode *--existing*: Prediction of a new batch of sequences using an existing mge-cluster model

For predicting the embedding coordinates and the cluster assignment of a new batch of plasmid sequences with an existing mge-cluster model, we designed the *--existing* operational mode. In this mode, mge-cluster requires an input file pointing to the nucleotide sequences of interest and the folder with the files constituting a mge-cluster model (Figure 5).

Mge-cluster performs the following steps: i) computes the same unitig definitions present in the file *\*unitigs*.*fasta*, using unitig-caller (--query mode), ii) uses the transform function from openTSNE python package to embed the new points to the existing embedding present in the file *\*embedding*.*pbz2*, iii) assigns the new points to the existing HDBSCAN clusters present in the file *\*clusters*.*pbz2* using the approximate_predict function from the hdbscan python package.

### Mge-cluster showcase: Generating an *E. coli* model to classify plasmid sequences

To showcase mge-cluster, we developed an *E. coli* model to classify plasmid sequences. We considered all plasmid sequences (n=6,864) with the species ‘*Escherichia coli’* annotated in the PLSDB database (20). Sequences from this database can contain near identical plasmids which could bias the downstream validation of mge-cluster. To select a single representative sequence among highly similar plasmids, we used cd-hit-est (version 4.8.1) to remove redundant sequences within a 0.99 sequence identity threshold (-c 0.99 -s 0.9 -aL 0.9) (36, 37). Cd-hit-est generated 6,185 groups encompassing plasmid sequences with high similarity and coverage, from these only a single representative sequence was chosen. The discarded sequences were used to benchmark the CPU time, runtime and memory required by mge-cluster to predict sequences considering an existing mge-cluster model. These sequences were also used as a quality check to ensure that mge-cluster returned the same cluster assignment as their cd-hit-est group.

We clustered the set of 6.185 non-redundant plasmids using mge-cluster. The perplexity was set to 100 (*--perplexity*), with a minimum cluster size of 30 (*--min_cluster*). Unitigs were discarded if their variance exceeded 0.01 (*--variance*). We used the script average_nucleotide_identity.py included in the pyani package (version 0.2.11) to calculate the average coverage and average nucleotide identity (ANI) of the plasmids within each cluster (38). We performed distinct runs of mge-cluster setting distinct perplexity values (10, 30, 50, 200) to compare the resulting clustering solution against the presented mge-cluster model (perplexity=100). For this, we considered the adjusted Rand index implemented in the function *adjustedRandIndex* from the mclust R package (version 5.4.7) (39). For representing the embedding created by openTSNE and the clusters defined by HDBSCAN, we used ggplot2 (version 3.3.6) and considered the Khachiyan algorithm implemented in the ggforce R package (40) to draw ellipses around the clusters.

The clustering given by mge-cluster was compared against the current typing schemes: i) ‘primary_cluster_id’ reported by the module MOB-typer of MOB-suite (12), a five-character fixed-length code that groups plasmids using complete-linkage clustering based on Mash distances (default distance = 0.06) and ii) plasmid taxonomic units (PTUs) reported by COPLA based on ANI distances and hierarchical stochastic block modelling (11). Due to the CPU time and memory required by COPLA to predict a single sample, we could not perform the typing and comparison of all the 6,185 plasmid sequences included in the model. Instead, from these 6,185 sequences, we considered 695 plasmids typed with a PTU in a recent publication introducing COPLA (10).

To quantify the concordance of the clustering solutions, we compared MOB-suite and COPLA against mge-cluster considering the adjusted Rand index (39). This metric compares two clustering solutions for the same set of points and returns a value ranging from 0 (no similarity) to 1 (identical clustering). The pairwise comparisons were only performed with sequences with a defined cluster for any of the typing tools, thus discarding plasmids labelled as -1 for mge-cluster or with an unknown PTU (‘-’) by COPLA. To further inspect the level of concordance between typing schemes, for each mge-cluster we computed its Simpson diversity for replicon, MOB-suite clusters (‘primary_cluster_id’) and COPLA PTUs. We considered the function *diversity* implemented in the vegan R package (version 2.5-7) specifying the ‘simpson’ index. This value can range from 0 (no diversity, same clustering solution) to 1 (high diversity, distinct clustering solution). To illustrate the differences between the clusterings given by mge-cluster and MOB-suite, we performed a gene synteny analysis with clinker (version v0.0.21) (41) using two randomly chosen sequences belonging to the same mge-cluster but differing in their MOB-suite cluster. To visualize the diversity of clustering solutions within each mge-cluster, we used the treemapify R package (version 2.5.5) which produces treemaps for displaying nested and hierarchical data (42).

To assess the performance of mge-cluster assigning plasmid sequences with a distinct gene content and origin, we considered all plasmid sequences (n=1,020) with the species ‘*Staphylococcus aureus’* annotated from the PLSDB database and used the operational mode --existing of mge-cluster to assign these sequences to the clusters defined in the *E. coli* mge-cluster model. In the same manner, we typed all IncN plasmids (n=206) from PLSDB belonging to a species different to *E. coli* annotated in the database and having uniquely a single replication gene in the field ‘PlasmidFinder’.

To illustrate the potential of mge-cluster to track the distribution of a gene-of-interest, we searched for AMR genes in our *E. coli* dataset using AMRFinderPlus (version 3.10.18) indicating as organism (-O) *Escherichia*, specifying the --plus flag and other default settings (43). From the resulting report, we searched for plasmid sequences encoding for the gene *mcr-1*.*1* (NCBI Reference Sequence accession NG_050417.1).

## Data and code availability

The mge-cluster package can be installed from bioconda https://anaconda.org/bioconda/mge-cluster under the open-source MIT license. Extensive documentation on mge-cluster usage is available at

https://gitlab.com/sirarredondo/mge-cluster.

The code required to reproduce the results and figures presented in this manuscript is available as a Rmarkdown document at

https://gitlab.com/sirarredondo/mge-cluster_manuscript

The plasmid sequences retrieved from the PLSDB database used to generate the *E. coli* mge-cluster for plasmid classification are publicly available at NCBI and their accession numbers listed on Supplementary Table S1. The accession numbers from the *S. aureus* and non-*E*.*coli* IncN plasmids retrieved from the PLSDB database and considered to assess the performance of mge-cluster typing new MGE data are available in Supplementary Tables S5 and S6 respectively.

The *E. coli* mge–cluster model presented in this manuscript is available as a figshare item at https://doi.org/10.6084/m9.figshare.21674078.v1

## Supplementary Figures

**Supplementary Figure S1.**
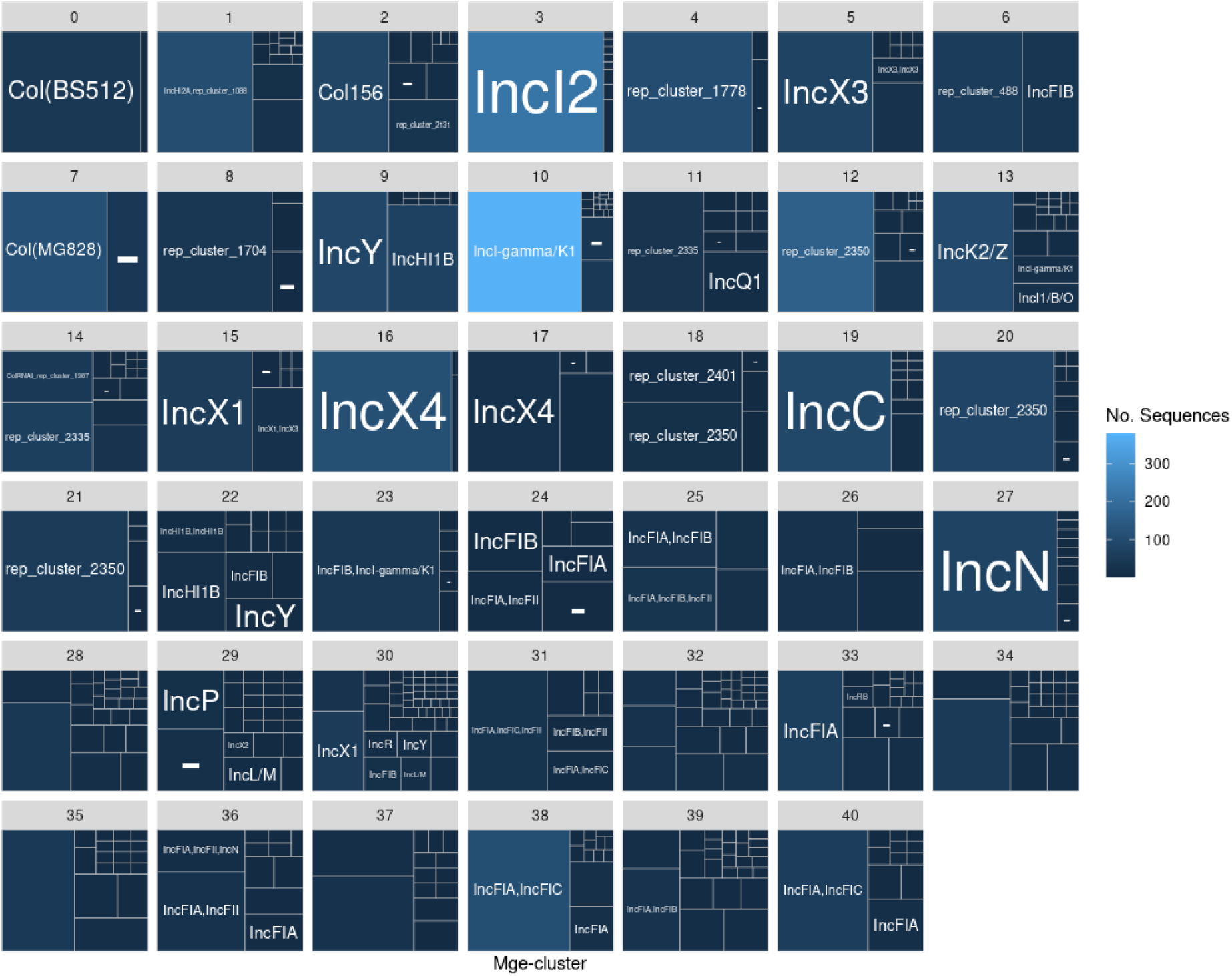
Replicon diversity reported by the module MOB-typer of MOB-suite in each mge-cluster (n=41). For each mge-cluster, the area of the plot is proportionally split into distinct tiles based on the number of plasmids with the same replicon combination. For each tile, the replicon combination is indicated in the center. In some cases, the tile may not contain any text because (i) the replicon combinations are rare resulting on a small tile where the text indicating the replicon(s) present cannot be fitted or (ii) multiple replicons are present in the plasmid resulting on a long text that surpasses the area of the tile.

**Supplementary Figure S2.**
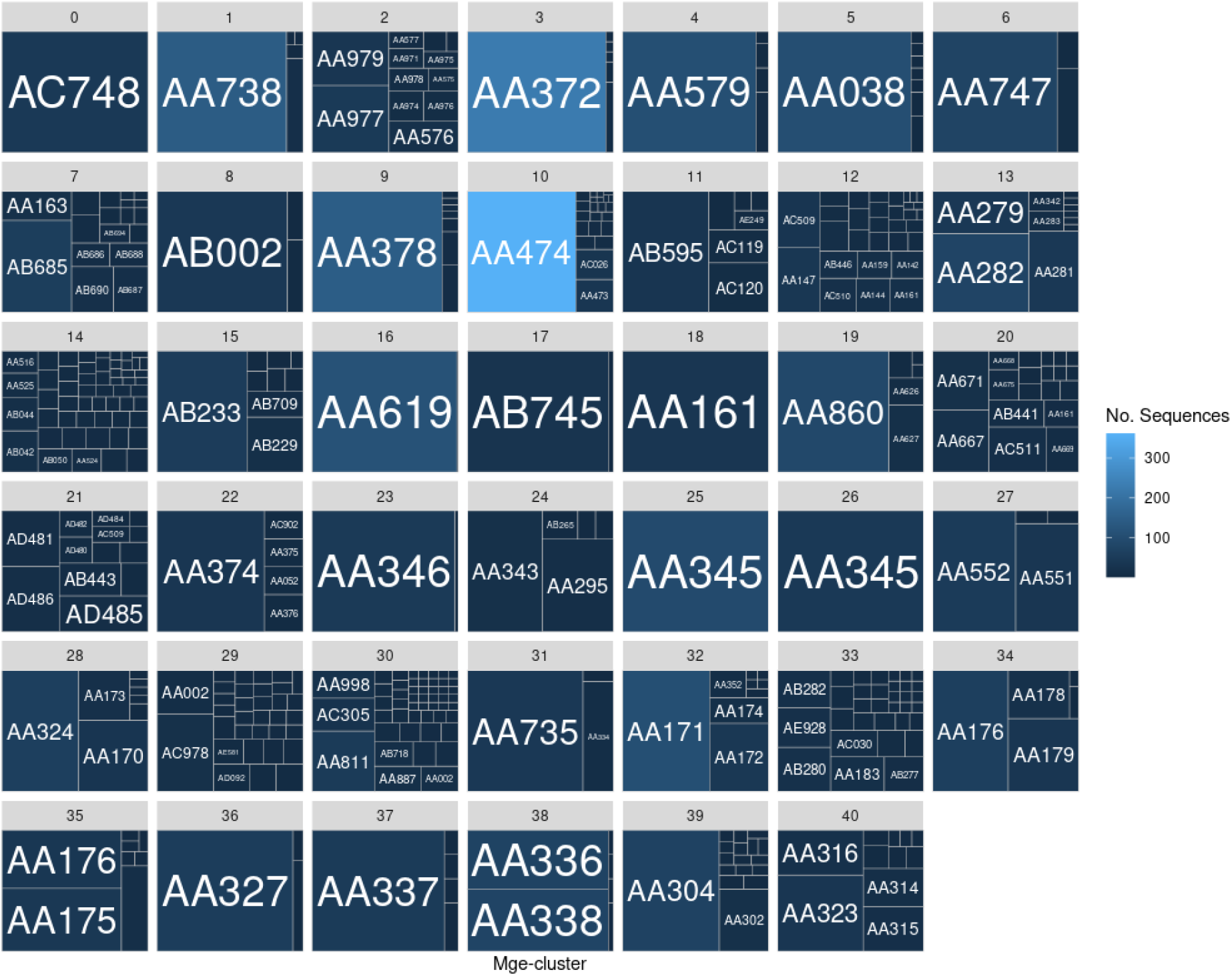
MOB-suite cluster diversity (‘primary_cluster_id’) present in each mge-cluster (n=41). For each mge-cluster, the area of the plot is proportionally split into distinct tiles based on the number of plasmids with the MOB-suite plasmid type. For each tile, the MOB-suite type is indicated in the center. In some cases, the tile may not contain any text because the MOB-suite type is rare among the mge-cluster resulting in a small area where the text cannot be fitted.

**Supplementary Figure S3.**
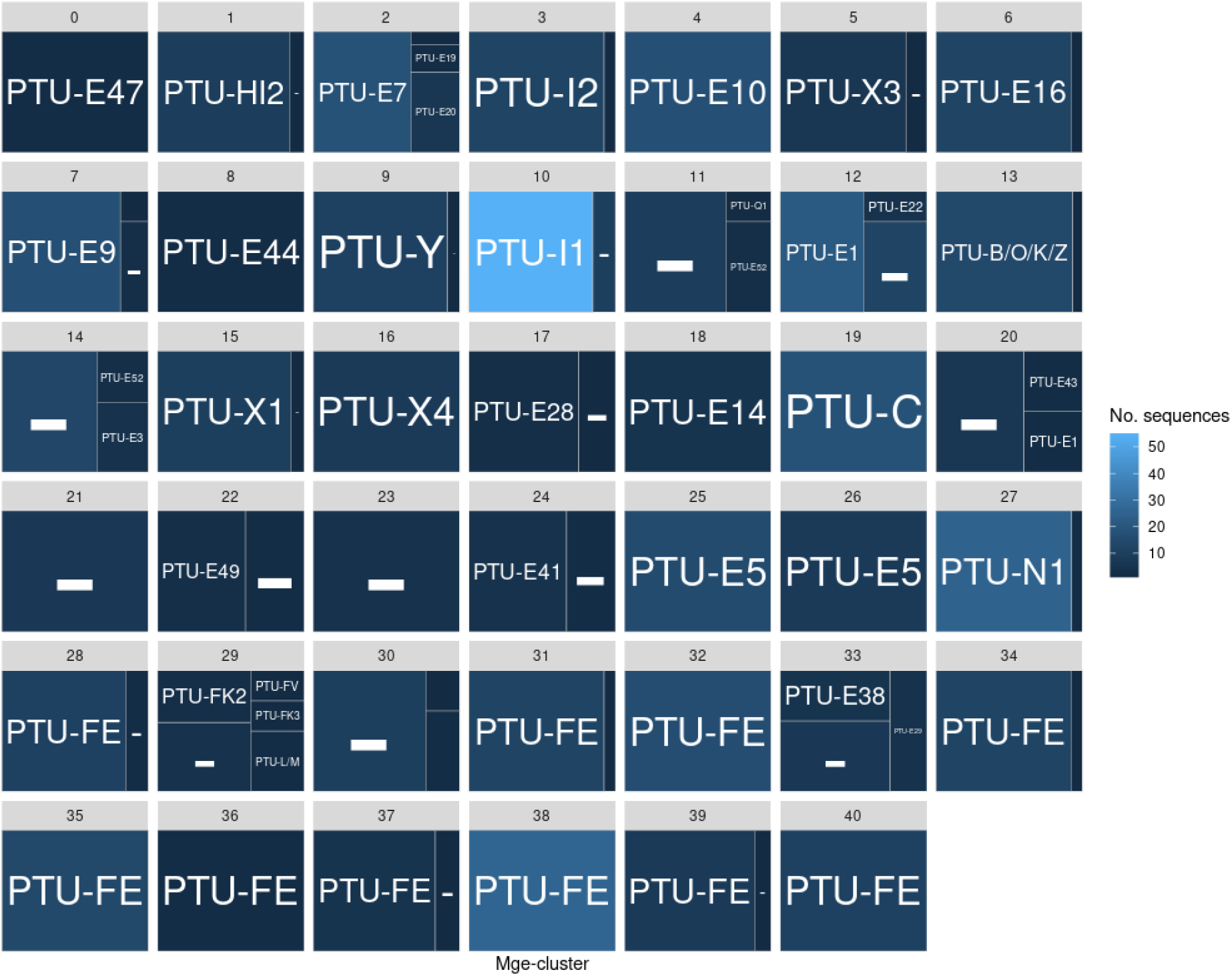
COPLA plasmid taxonomic unit (PTU) diversity present in each mge-cluster. For each mge-cluster, the area of the plot is proportionally split into distinct tiles based on the number of plasmids with a particular COPLA PTU. This diversity is uniquely based on 695 plasmid sequences previously typed in COPLA’s original publication (10). In some cases, COPLA labelled sequences as ‘-’ corresponding to plasmids with an unknown PTU.

**Supplementary Figure S4.**
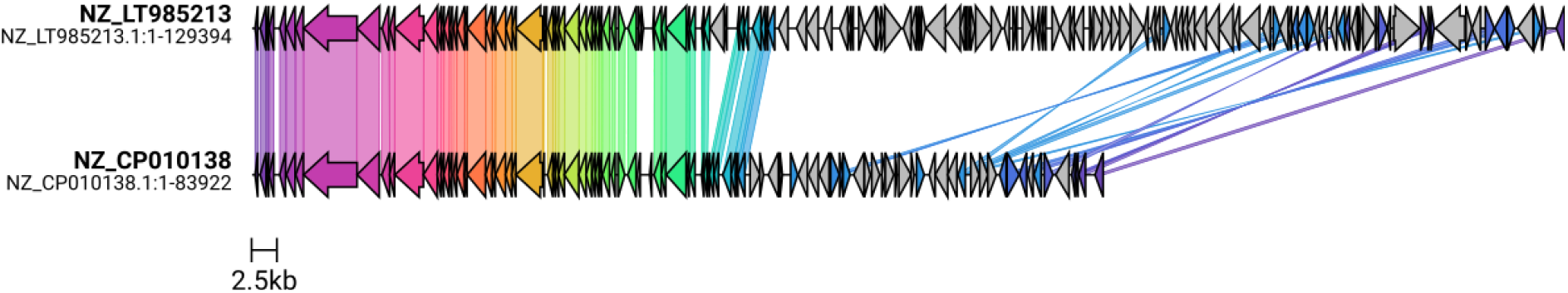
Gene synteny plot between the sequences NZ_LT985213.1 (top) NZ_CP010138.1 (bottom) belonging to mge-cluster 31. These two sequences were randomly picked and represented the two major types (AA735 and AA334, respectively) defined by MOB-suite. The plot was created with clinker (41) based on the genome annotation (.gbk) computed with prokka (44), and homologous genes with a minimum identity of 80% are indicated with a link between the two sequences.

**Supplementary Figure S5.**
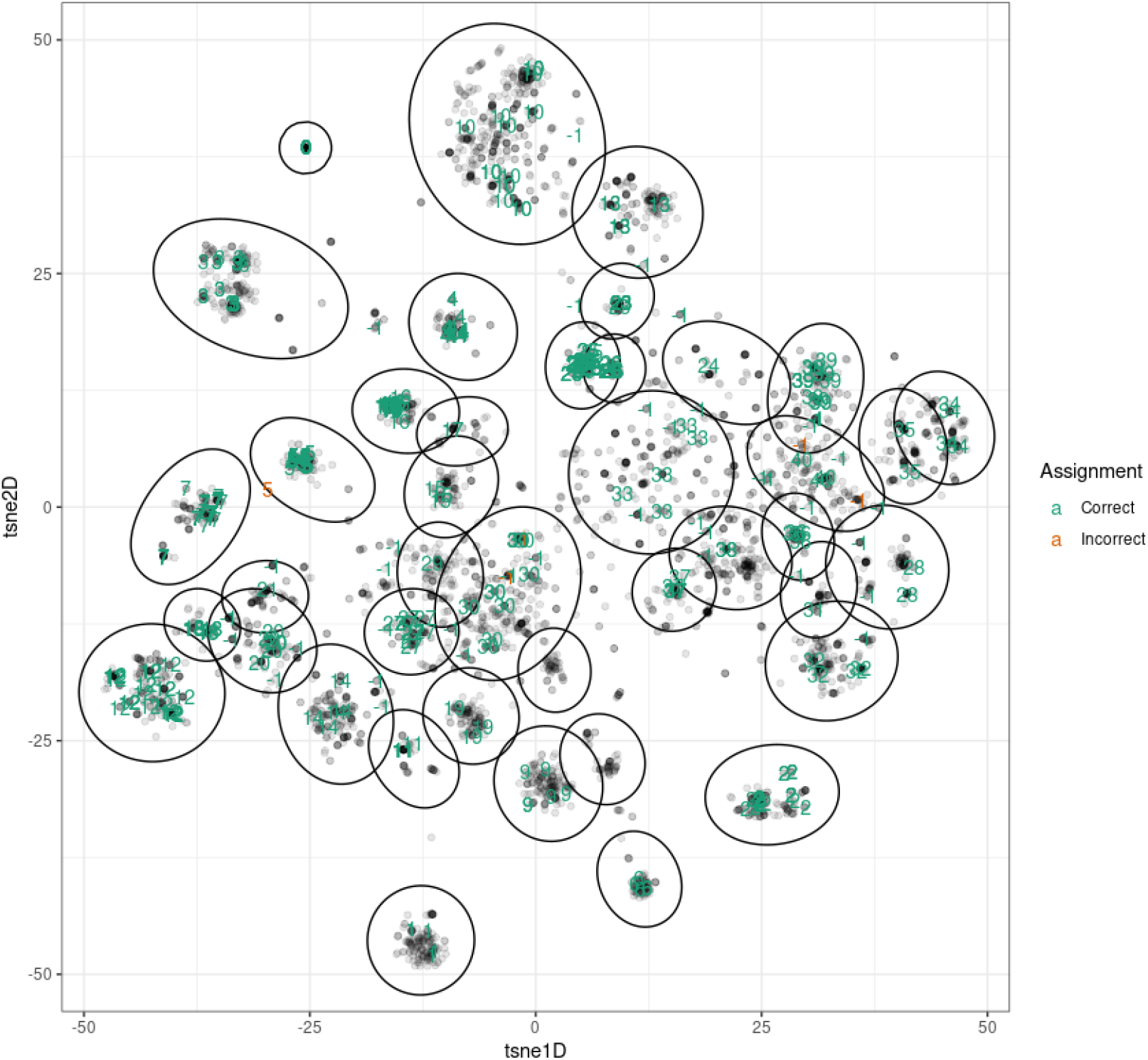
Embedding and assignment of the plasmid sequences (n=675) that were originally discarded by cd-hist-est, and considered as a benchmarking set. These sequences are labelled based on their predicted HDBSCAN cluster and coloured based on whether their assignment was correct (in green) or incorrect (in orange).

